# Purification of recombinant SARS-CoV-2 spike, its receptor binding domain, and CR3022 mAb for serological assay

**DOI:** 10.1101/2020.07.31.231282

**Authors:** Kang Lan Tee, Philip J. Jackson, Joseph M. Scarrott, Stephen R. P. Jaffe, Abayomi O. Johnson, Yusuf Johari, Thilo H. Pohle, Theo Mozzanino, Joseph Price, James Grinham, Adam Brown, Martin J. Nicklin, David C. James, Mark J. Dickman, Tuck Seng Wong

## Abstract

Serology testing for COVID-19 is highly attractive because of the relatively short diagnosis time and the ability to test for an active immune response against the SARS-CoV-2. In many types of serology tests, the sensitivity and the specificity are directly influenced by the quality of the antigens manufactured. Protein purification of these recombinantly expressed viral antigens [*e.g.*, spike and its receptor binding domain (RBD)] is an important step in the manufacturing process. Simple and high-capacity protein purification schemes for spike, RBD, and CR3022 mAb, recombinantly expressed in CHO and HEK293 cells, are reported in this article. The schemes consist of an affinity chromatography step and a desalting step. Purified proteins were validated in ELISA-based serological tests. Interestingly, extracellular matrix proteins [most notably heparan sulfate proteoglycan (HSPG)] were co-purified from spike-expressing CHO culture with a long cultivation time. HSPG-spike interaction could play a functional role in the pathology and the pathogenesis of SARS-CoV-2 and other coronaviruses.

## INTRODUCTION

Severe acute respiratory syndrome coronavirus 2 (SARS-CoV-2), identified in December 2019, is the causative agent for the global pandemic designated COVID-19 [1]. Serological surveys represent an invaluable tool to study immune response, to assess the extent of the pandemic given the existence of asymptomatic cases and to guide control measures. There is clear research evidence that SARS-CoV-2 spike (S) and nucleocapsid (N) proteins are the primary viral antigens, against which antibodies are raised [2]. Hence, ELISA-based serological tests rely on the purified SARS-CoV-2 components that are recombinantly expressed in insect or mammalian cells.

Coronavirus entry into host cells is mediated by the transmembrane spike glycoprotein that forms homotrimers protruding from the viral surface [3]. The spike protein contains a receptor binding domain (RBD) that specifically recognizes angiotensin-converting enzyme 2 (ACE2) as its receptor [4, 5]. Spike is therefore a prime target of neutralising antibody and vaccine [6].

CR3022 mAb, isolated previously from a convalescent SARS patient, is a neutralizing antibody that targets the RBD of SARS-CoV [7]. The spike proteins of SARS-CoV-2 and SARS-CoV are phylogenetically closely related, sharing an amino acid sequence identity of ∼77% [8]. Not surprising, CR3022 also binds to the RBD of SARS-CoV-2, as confirmed by ELISA [9], bio-layer interferometry (BLI; [9]) and protein crystallography [10]. For the purpose of a serological test, CR3022 serves as a good positive control, when either spike or RBD is used as an antigen.

The focus of this article is two-fold. First, we present detailed protein purification schemes for spike, RBD and CR3022, three important components for an ELISA-based serological test. These schemes, often omitted from or briefly described in publications, are optimised specifically for the ÄKTA protein purification system to increase protein manufacturing capacity. Second, we report and discuss co-purified protein impurities, which could have significant implications on the downstream application of these recombinant proteins and analysis of their associated experimental data.

## METHODS

### Recombinant SARS-CoV-2 spike, RBD, and CR3022 mAb

#### Transient spike/RBD production in HEK

Expi293F cells (Thermo Fisher Scientific) were routinely cultivated in Expi293F Expression medium (Thermo Fisher Scientific) at 37°C, 125 rpm in 8% CO_2_, ∼80% humidity. Cells were routinely sub-cultured every 3 – 4 days, by seeding at 0.3 – 0.4 0.4 mvc/mL. Cells were transfected with plasmid DNA [11, 12] using PEI under optimized conditions, and cultured for up to 8 days.

#### Stable spike production in CHO

A CHO-S derived clone was used to stably express spike. Cultures were grown in CD-CHO (Thermo Fisher Scientific) under biphasic conditions in an orbital shaker at 37°C, 140 rpm, 5% CO_2_, 85% humidity, with a feeding of 5% (v/v) CHO CD EfficientFeed B (Thermo Fisher Scientific) on days 2, 4, 6 and 8 of a 10-day fed batch cultivation.

#### Stable CR3022 production in CHO

A CHOK1SV GS-KO cell line was used to stably express CR3022. Cultures were grown in CD-CHO (Thermo Fisher Scientific) in an orbital shaker at 37°C, 140 rpm, 5% CO_2_, 85% humidity, with a feeding of 10% (v/v) CHO CD EfficientFeed B (Thermo Fisher Scientific) on days 3, 6 and 9 of an 11-day fed batch cultivation.

### Protein purification of SARS-CoV-2 spike and RBD

All protein purifications in this study was conducted using an ÄKTA Pure system (Cytiva). A 5-mL HisTrap™ HP column (Cytiva) was washed with 5 column volumes (CVs) of buffer B (50 mM sodium phosphate, 300 mM NaCl, 250 mM imidazole, pH 8.0), and equilibrated with 5 CVs of buffer A (50 mM sodium phosphate, 300 mM NaCl, 10 mM imidazole, pH 8.0). Spent medium containing either spike or RBD was filtered using a 0.22 μm stericup and loaded onto the equilibrated column using a sample pump. Typically, a volume of 50 – 500 mL was loaded. After sample loading, the column was washed in three steps using 5 CVs of buffer A, 5 CVs of 4.5% (v/v) buffer B, and 10 CVs of 9% (v/v) buffer B. Protein was eluted using 100% (v/v) buffer B, and collected in fractions (1 mL/fraction) using a fraction collector. The column equilibration, sample loading, and column washing steps were conducted at 5 mL/min, while the elution step at 1 mL/min. To further reduce non-specific binding, spent medium was adjusted to 20 mM imidazole using buffer B, prior to filtration and sample loading.

### SDS-PAGE

Protein fractions were analysed on a NuPAGE™ 4-12%, Bis-Tris, 1 mm, 12-well mini protein gel (Thermo Fisher Scientific) to check for size, purity, and integrity. The gel was run using NuPAGE™ MES SDS running buffer (Thermo Fisher Scientific) at a constant voltage of 200 V for 40 min. For visualization, the gel was stained using InstantBlue™ (Expedeon).

### Buffer exchange and storage of SARS-CoV-2 spike and RBD

Protein fractions were pooled and buffer exchanged into storage buffer [20 mM Tris, 200 mM NaCl, 10% (v/v) glycerol, pH 8.0] using a PD-10 desalting column (Cytiva). Protein sample was aliquoted, snap frozen in liquid nitrogen, and stored at −80°C.

### Protein purification of CR3022 mAb

A 5-mL HiTrap™ Protein G HP column (Cytiva) was equilibrated with 10 CVs of Protein G IgG Binding Buffer (pH 5.0; Thermo Fisher Scientific). Spent medium containing CR3022 mAb was diluted with binding buffer at a 1:1 volume ratio and filtered using a 0.22 μm stericup, prior to sample loading using a sample pump. The column was washed with 10 CVs of binding buffer. mAb was eluted using IgG Elution Buffer (pH 2.8; Thermo Fisher Scientific). Protein fractions (1 mL/fraction) were collected using a fraction collector and collection tubes containing 100 μL/tube of neutralization buffer (1 M Tris, pH 9.0). The column equilibration, sample loading, and column washing steps were conducted at 5 mL/min, while the elution step at 1 mL/min.

### Buffer exchange and storage of CR3022 mAb

After SDS-PAGE analysis, mAb fractions were pooled and buffer exchanged into storage buffer [phosphate-buffered saline, 50% (v/v) glycerol, 0.02% (v/v) ProClin 300] using a PD-10 desalting column (Cytiva). mAb sample was aliquoted and stored at −20°C.

### Protein quantification

Spike, RBD, and CR3022 mAb samples were diluted with their corresponding storage buffers, and quantified using Pierce™ Coomassie Plus (Bradford) Assay Kit (Thermo Fisher Scientific). Bovine serum albumin (2.5 μg/mL to 25 μg/mL) was used as a protein calibration standard. Briefly, 100 μL of Bradford reagent was added to 100 μL of diluted protein sample in a 96-well microplate. After a 30-sec shaking, the plate was incubated at room temperature for 10 min. Absorbance was then measured at 595 nm using a Multiskan™ FC microplate photometer (Thermo Fisher Scientific).

### Western biot

Proteins were resolved on a NuPAGE™ 4-12%, Bis-Tris, 1 mm, 12-well mini protein gel (Thermo Fisher Scientific), and transferred onto a PDVF membrane (iBlot™ 2 Transfer Stacks; Thermo Fisher Scientific). His-tagged protein was detected using a mouse anti-6×His IgG conjugated to horseradish peroxidase (MCA1396P; Bio-Rad), and the blot was developed using Pierce™ ECL Western Blotting Substrate (Thermo Fisher Scientific).

### Peptide analysis by mass spectrometry

The identification of proteins analysed by SDS-PAGE was performed by first excising the bands from the gel and slicing them into 1 mm pieces. The gel pieces were then de-stained with 2 x 200 μL of 50% (v/v) acetonitrile, 50 mM ammonium bicarbonate (ACN/ABC). The proteins were reduced and S-alkylated with 10 mM TCEP and 55 mM MMTS, respectively, with washing in ACN/ABC after each reaction. The gel pieces were dried in a vacuum centrifuge then rehydrated in 50 mM ABC containing 12.5 ng/μL trypsin (Promega), 5 mM calcium chloride. Proteolytic digestion was by incubation at 37°C for 16 h. Peptides were extracted from the gel pieces with 15 μL of 25 mM ABC then 20 μL of neat ACN. The extracts were combined, dried in a vacuum centrifuge and the peptides redissolved in 0.5% (v/v) TFA, 3% (v/v) ACN for analysis by liquid chromatography coupled to mass spectrometry (LC-MS/MS) using an RSLCnano-Q Exactive HF system (Thermo Fisher Scientific). Mass spectra were searched against Chinese hamster or human proteome databases (with inserted Spike and RBD sequences) using Mascot software (Matrix Science).

For identification of proteins in solution, samples were concentrated to approximately 1 g/L by centrifugal ultrafiltration and 5 μg diluted to 7 μL with water. 1 μL each of 500 mM ABC, 0.2% (w/v) ProteaseMax surfactant (Promega) in 50 mM ABC and 0.2 g/L trypsin was then added. The proteins were incubated at 37°C for 16 h then 1 μL of 5% (v/v) TFA was added to degrade the surfactant. After incubation at RT for 5 min, the peptides were desalted using a C18 spin column (Thermo Fisher Scientific) then dried and redissolved for LC-MS/MS analysis as above.

### Size exclusion chromatography

A HiLoad™ 26/600 Superdex™ 200 pg column (Cytiva) was washed with 2 CVs of water, and equilibrated with 1.5 CVs of buffer (20 mM Tris, 200 mM NaCl, pH 8.0). A 2-mL protein sample was injected via a 5-mL sample loop. Separation was performed at a flow rate of 2 mL/min. Eluate was collected in fractions (4 mL/fraction) using a fraction collector.

### Co-immunoprecipitation

Co-immunoprecipitation was conducted using a NAb™ Spin Kit (Protein G, 0.2 mL; Thermo Fisher Scientific). Briefly, the columns were equilibrated with 400 μL of binding buffer (100 mM phosphate, 150 mM NaCl, pH 7.2), and incubated with 500 μL of protein sample for 10 min at room temperature. The columns were subsequently washed 3 times with 400 μL of binding buffer, followed by 3 elution steps with 400 μL of IgG Elution Buffer. Eluates were collected in tubes containing 40 μL/tube of neutralization buffer (1M Tris-HCl, pH 8.5).

## RESULTS & DISCUSSION

### SARS-CoV-2 spike and RBD constructs

The gene sequence for spike, in the constructs provided by Krammer [11, 12], was derived from the genomic sequence of the first virus isolate, designated as Wuhan-Hu-1 (GenBank: MN908947.3; Fig. 1) [13]. The spike protein sequence contains multiple modifications intended for protein stabilization and downstream processing: (a) the polybasic cleavage site (R682RAR685) recognised by a furin [14] was removed via an alanine replacement, (b) two stabilizing mutations (K986P and V987P; [15]) were introduced, (c) the transmembrane domain (a.a. 1214 – 1236) and the endodomain (a.a. 1237 – 1273) were truncated, and (d) a thrombin cleavage site (LVPR↓GS), a trimerization domain from the bacteriophage T4 fibritin (GYIPEAPRDGQAYVRKDGEWVLLSTFL) [16, 17], and a hexahistidine tag (HHHHHH) were fused to P1213. For the RBD construct, the native N-terminal signal peptide of spike (MFVFLVLLPLVSSQ) was fused to the RBD sequence (a.a. 319 – 541), and joined with a C-terminal hexahistidine tag.

**Figure 1:**
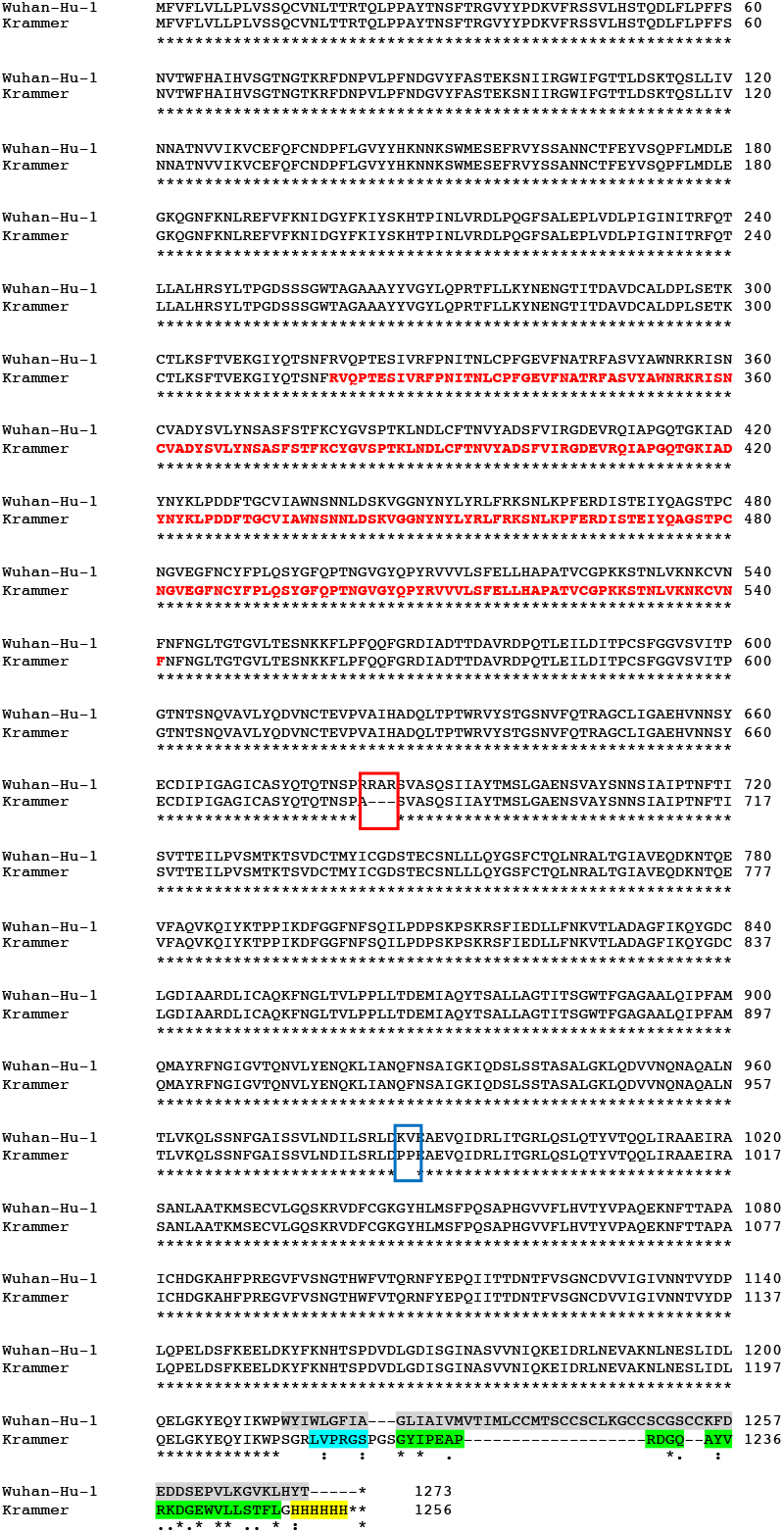
Sequence alignment of the S protein from the severe acute respiratory syndrome coronavirus 2 isolate Wuhan-Hu-1 (GenBank: MN908947.3) and the S protein variant used in this study (provided by Krammer), created using Clustal Omega [35]. In the S variant, the polybasic cleavage site (R682RAR685) recognised by a furin was replaced by an alanine (boxed in red). Two stabilizing mutations (K986P and V987P) were introduced (boxed in blue). The transmembrane domain (a.a. 1214 – 1236) and the endodomain (a.a. 1237 – 1273), highlighted in grey, were truncated. A thrombin cleavage site (cyan), a trimerization domain from the bacteriophage T4 fibritin (green) and a hexahistidine tag (yellow) were fused to P1213. For the RBD construct used in this study, the native N-terminal signal peptide of SARS-CoV-2 S protein (MFVFLVLLPLVSSQ) was fused to the RBD sequence (a.a. 319 – 541; sequence in bold red), and joined with a C-terminal hexahistidine tag.

### Protein purification of spike

All the purification schemes in this study were developed for use with ÄKTA protein purification systems and the likes. The schemes, consisting of an affinity chromatography step and a desalting step, were designed for routine larger-scale preparative purification. The number of purification steps was intentionally minimised to strike a balance between protein purity that suffices a serological test and protein yield, and to reduce processing time.

SDS-PAGE analysis showed that the full-length spikes purified from both HEK and CHO cultures were relatively pure and homogenous (Fig. 2A), giving a Mw of >180 kDa that is significantly larger than the calculated M_w_ of 139.2 kDa. SARS-CoV-2 S gene encodes 22 N-linked glycan sequons (N-X-S/T, where X ≠ P) per protomer [18]. The size difference of ∼40 kDa indicated that the recombinant spike is a heavily glycosylated protein. The band corresponding to the spike protein was subsequently analysed using in-gel trypsin digestion in conjunction with mass spectrometry analysis. Mass spectrometric data confirmed the identity of the spike (Fig. 2B). Contrary to the work reported by Krammer and co-workers, in which the full-length spike appeared as a double band (∼180 kDa and ∼130 kDa) on a reducing SDS-PAGE which was postulated due to protein degradation resulting in a smaller protein [11], our purified spike ran as a clear single band on both non-reducing (Fig. 2A) and reducing (data not shown) SDS-PAGE.

**Figure 2:**
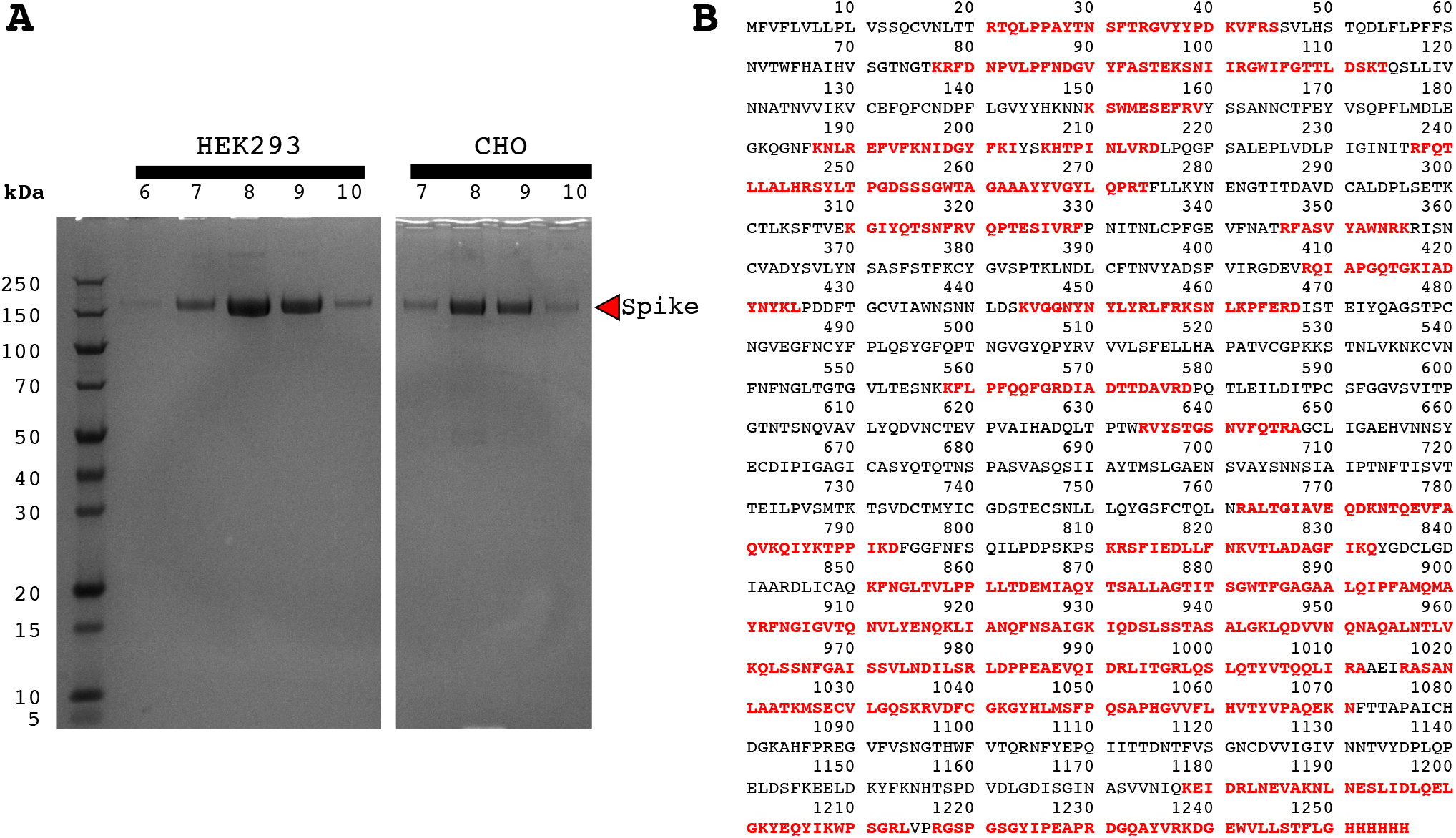
**(A)** Fractions collected from a step elution in SARS-CoV-2 spike purification using an affinity chromatography (5-mL HisTrap™ HP column). **(B)** Gel band corresponding to spike was excised from the SDS-PAGE, post visualisation by InstantBlue™. The gel band was de-stained and proteolytically digested with trypsin to generate peptides for mass spectrometric analysis using LC-MS/MS. Peptides mapped to spike protein sequence were coloured in red.

### Protein purification of RBD

The same purification scheme, developed for the full-length spike, was applied for the purification of RBD. Non-reducing SDS-PAGE analysis of the purified RBD showed two distinct bands (Fig. 3), a dominant band with a Mw of ∼30 kDa and a side band with a Mw of ∼60 kDa. As our RBD sequence (a.a. 319 – 541) contains, in total, 9 cysteine residues (C336, C361, C379, C391, C432, C480, C488, C525 and C538), we suspected an RBD dimer formation. When we re-ran the purified RBD on a reducing SDS-PAGE, a clear single band was obtained (Fig. 3), confirming our hypothesis. The bands corresponding to the RBD dimer and RBD monomer were analysed using in-gel trypsin digestion in conjunction with mass spectrometry analysis and confirmed the identity of RBD in both cases (Fig. 4). Based on the crystallographic data of RBD (6M0J, [19]), there are four disulphide bonds in RBD (C336-C361, C379-C432, C391-C525 and C480-C488; Fig. 5). The free C538 is likely the cysteine residue involved in inter-molecular dimerization.

**Figure 3:**
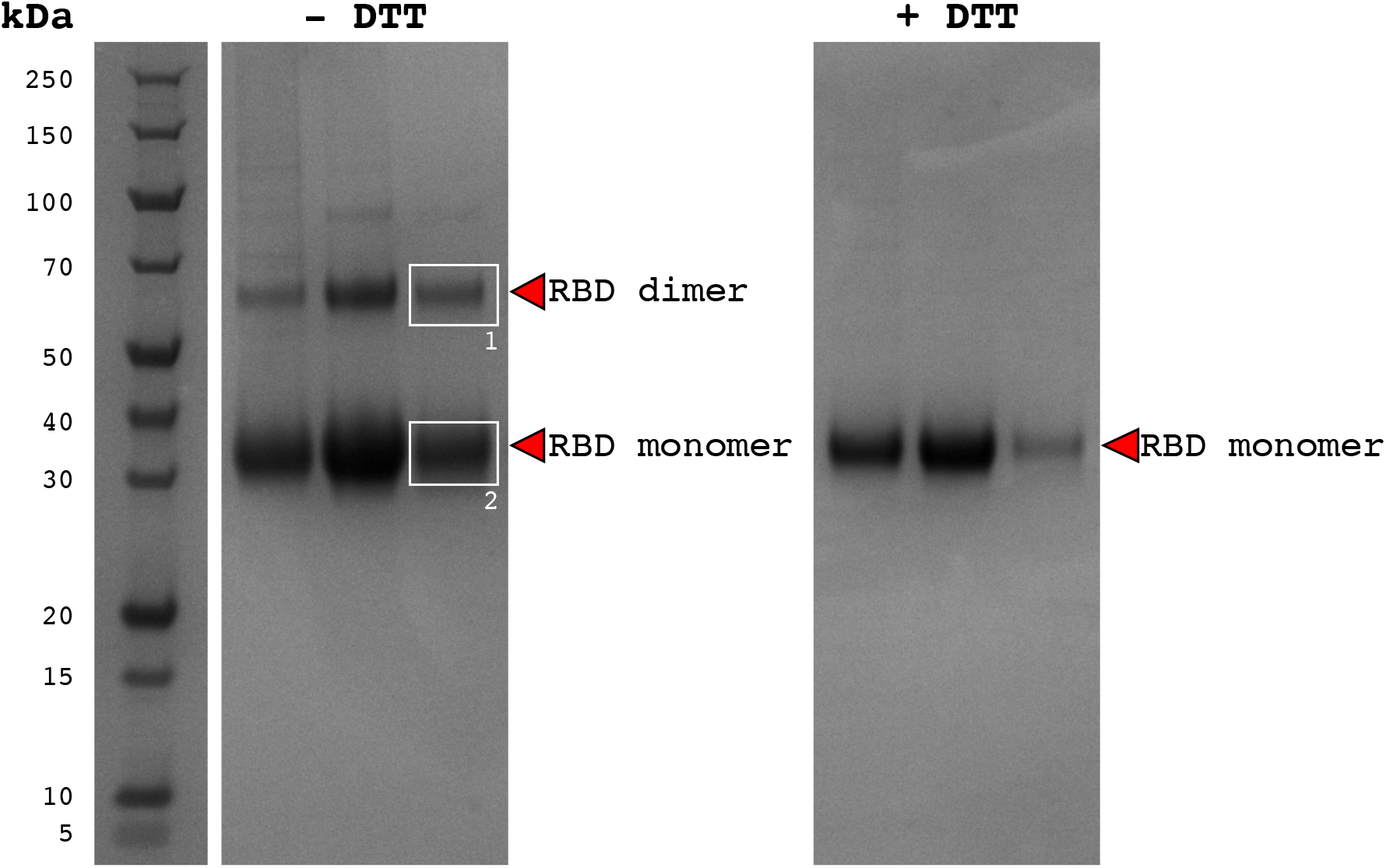
Purified receptor binding domain (RBD; a.a. 319 – 541) of SARS-CoV-2 spike was analysed on a non-reducing *(left)* and reducing *(right)* SDS-PAGE, confirming the RBD’s dimerization.

**Figure 4:**
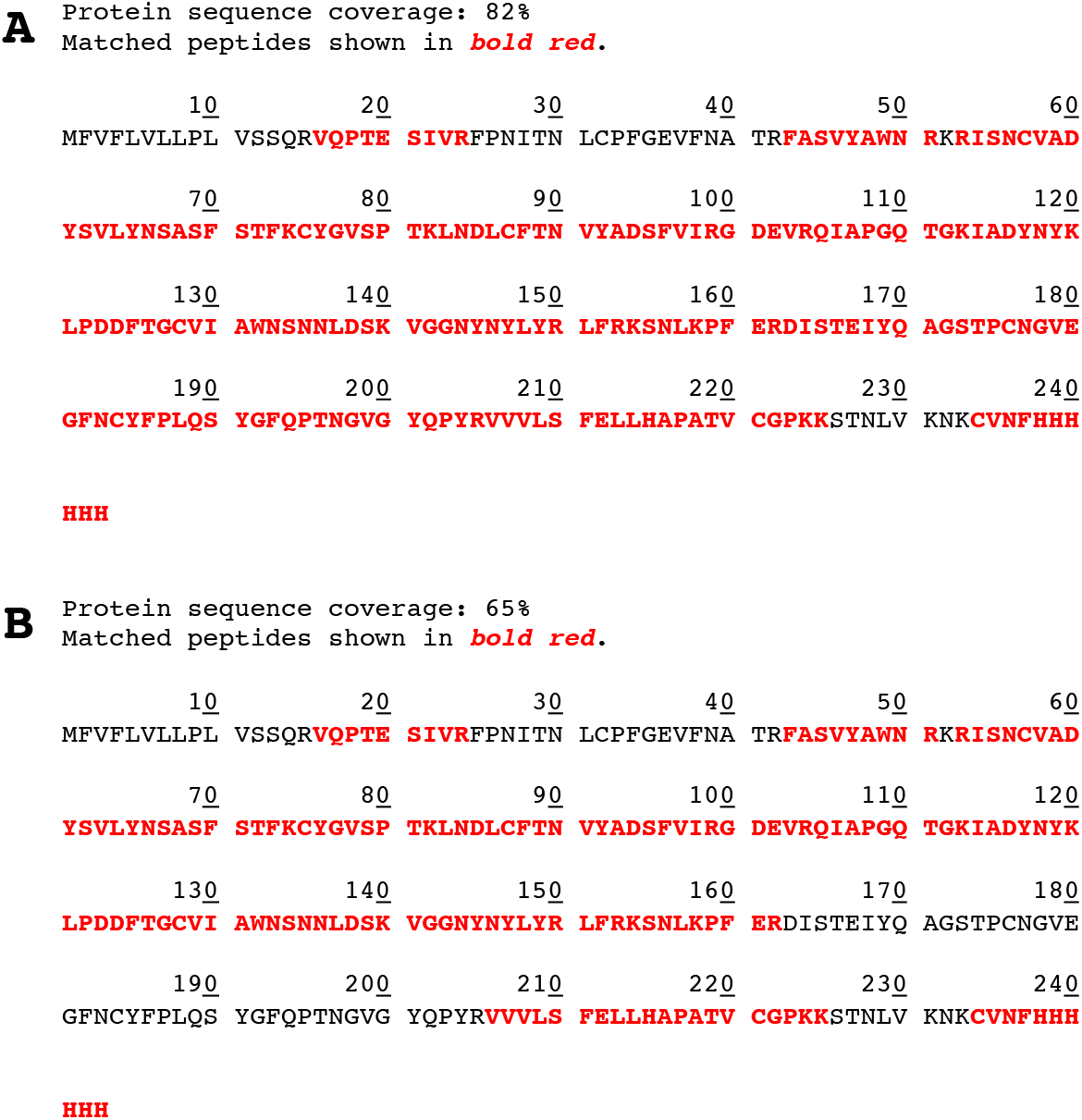
The boxed gel bands in Fig. 3 (box 1 for RBD dimer and box 2 for RBD monomer) were excised from the SDS-PAGE, post visualisation by InstantBlue™. The gel bands were de-stained and proteolytically digested with trypsin to generate peptides for mass spectrometric analysis using LC-MS/MS. Peptides mapped to RBD sequence were coloured in red: **(A)** Box 1 – RBD dimer, and **(B)** Box 2 – RBD monomer.

**Figure 5:**
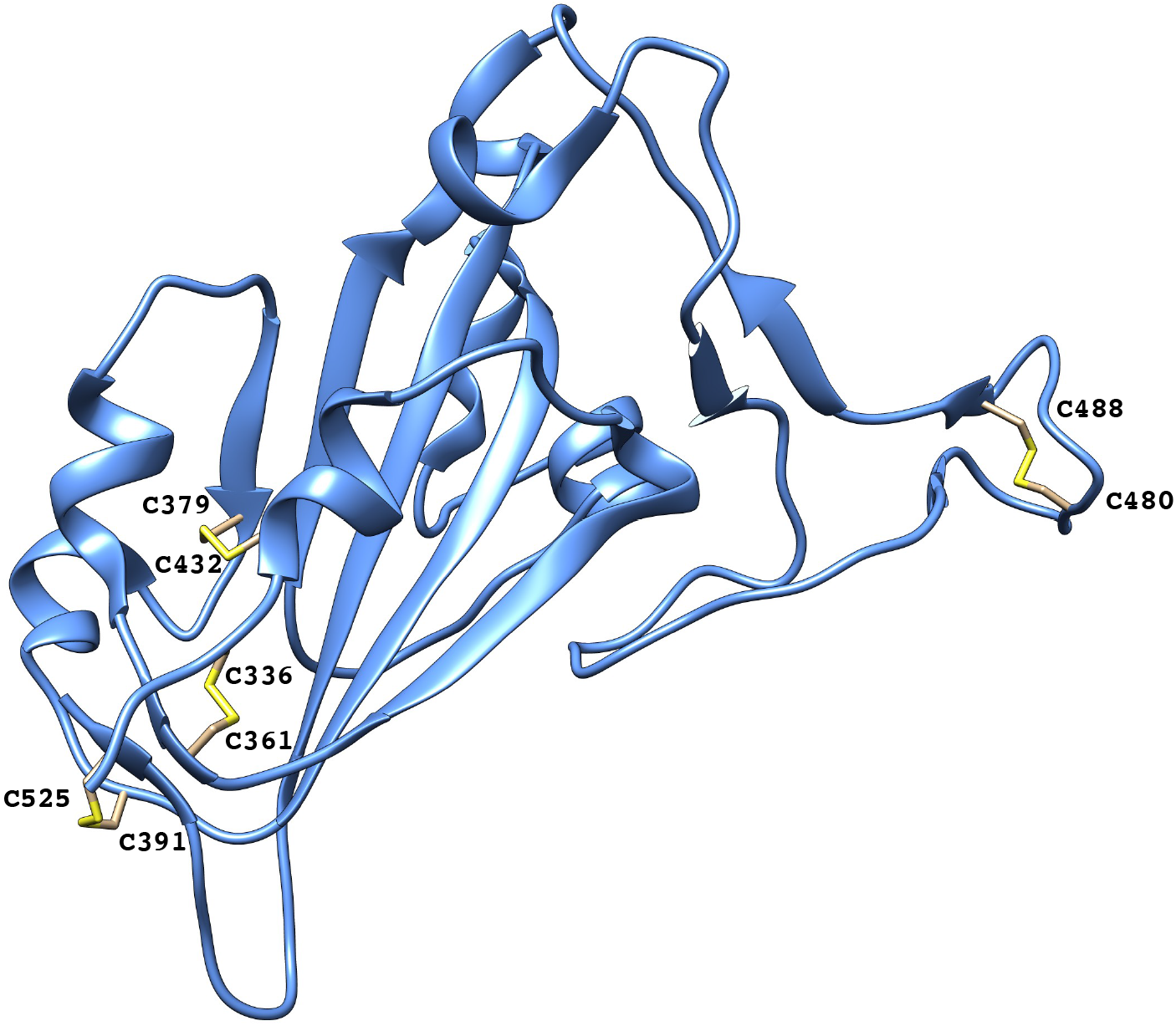
Ribbon diagram of RBD structure (PBD 6M0J), created using CHIMERA [36]. The four disulphide bonds in RBD (C336-C361, C379-C432, C391-C525 and C480-C488) were shown in sticks.

Like the recombinant full-length spike, recombinant RBD also appeared larger in size. There are two N-glycosylation sites within the RBD sequence (N331 and N343) [18]. This could explain the size difference of ∼4 kDa between the Mw deduced from the SDS-PAGE (∼30 kDa) and the calculated Mw (26.1 kDa).

### Protein purification of CR3022 mAb

CR3022 mAb was purified using a pre-packed protein G column, with a straightforward purification scheme. This IgG interacts strongly with protein G, evidenced by its broad elution peak spanning over 16 fractions (Fig. 6). We chose a Protein G IgG Binding Buffer with a pH of 5.0 in our purification scheme for a stronger IgG interaction with protein G. The binding between protein G and IgG was shown to be pH-dependent between 2.8 and 10, strongest at pH 4 and 5, and weakest at pH 10 [20]. The downside of using an acidic binding buffer is protein precipitation, which occurred when the spent medium was diluted with the binding buffer at a 1:1 volume ratio. The yield of CR3022 mAb (57 mg/L spent medium) was similar to that estimated from HPLC using MAbPac™ Protein A column (40 – 100 mg/L spent medium). The precipitated protein was likely host cell proteins. It is therefore necessary to filter the diluted spent medium with a 0.22 μm stericup, prior to sample loading onto the equilibrated protein G column. Non-reducing (Fig. 7A) and reducing (Fig. 7B) SDS-PAGE analyses confirmed the presence of both heavy chain and light chain in the purified CR3022.

**Figure 6:**
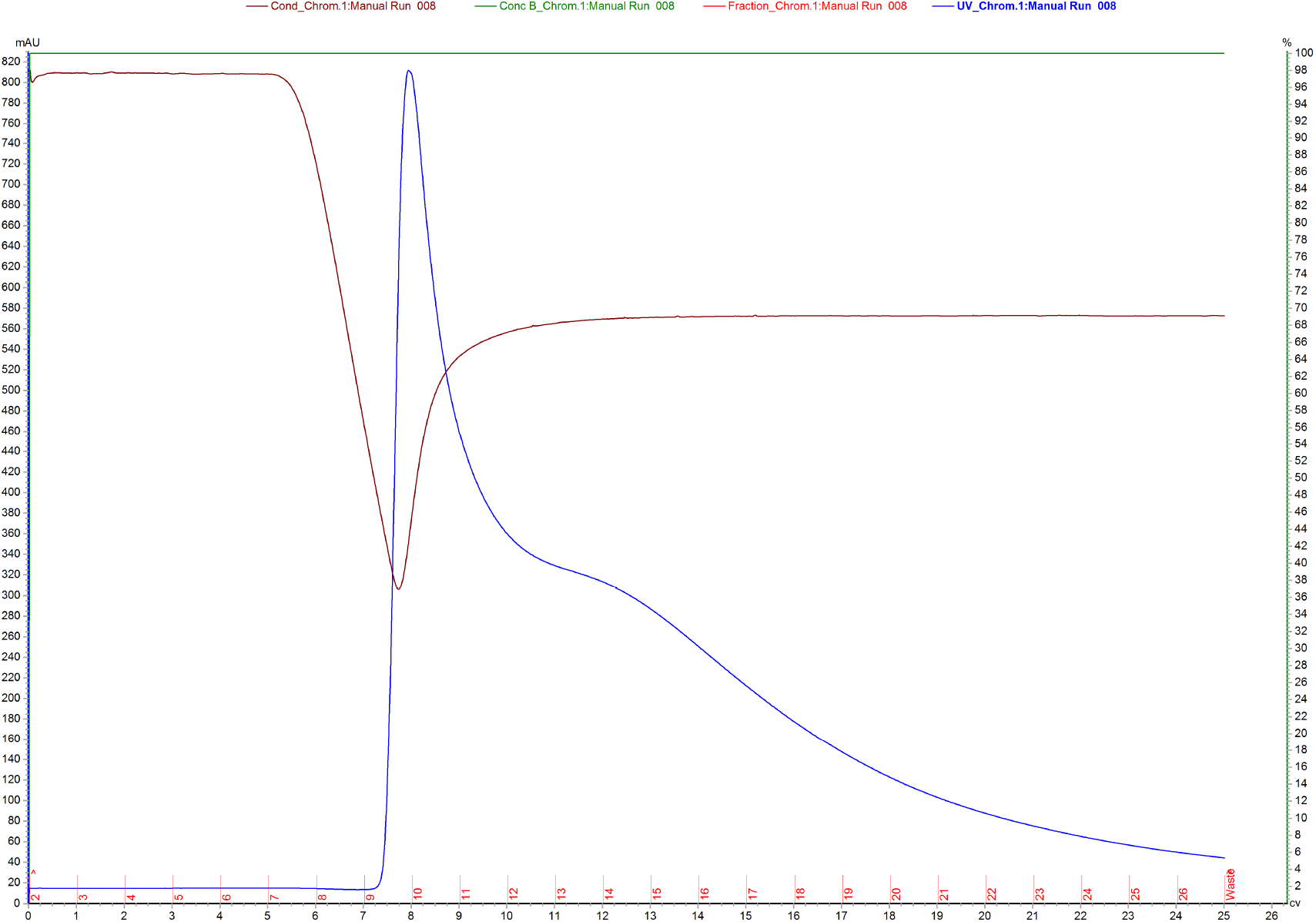
The chromatogram of a step elution in CR3022 mAb purification using affinity chromatography (5-mL HiTrap™ Protein G HP column). The x axis, left y axis, and right y axis show the elution volume in mL, UV absorbance at 280 nm, and % (v/v) of buffer B, respectively. The blue, green and brown curves represent UV absorbance at 280 nm, % (v/v) of buffer B and conductivity, respectively. CR3022 mAb interacted strongly with Protein G, and was eluted in a broad peak spanning over 16 fractions.

**Figure 7:**
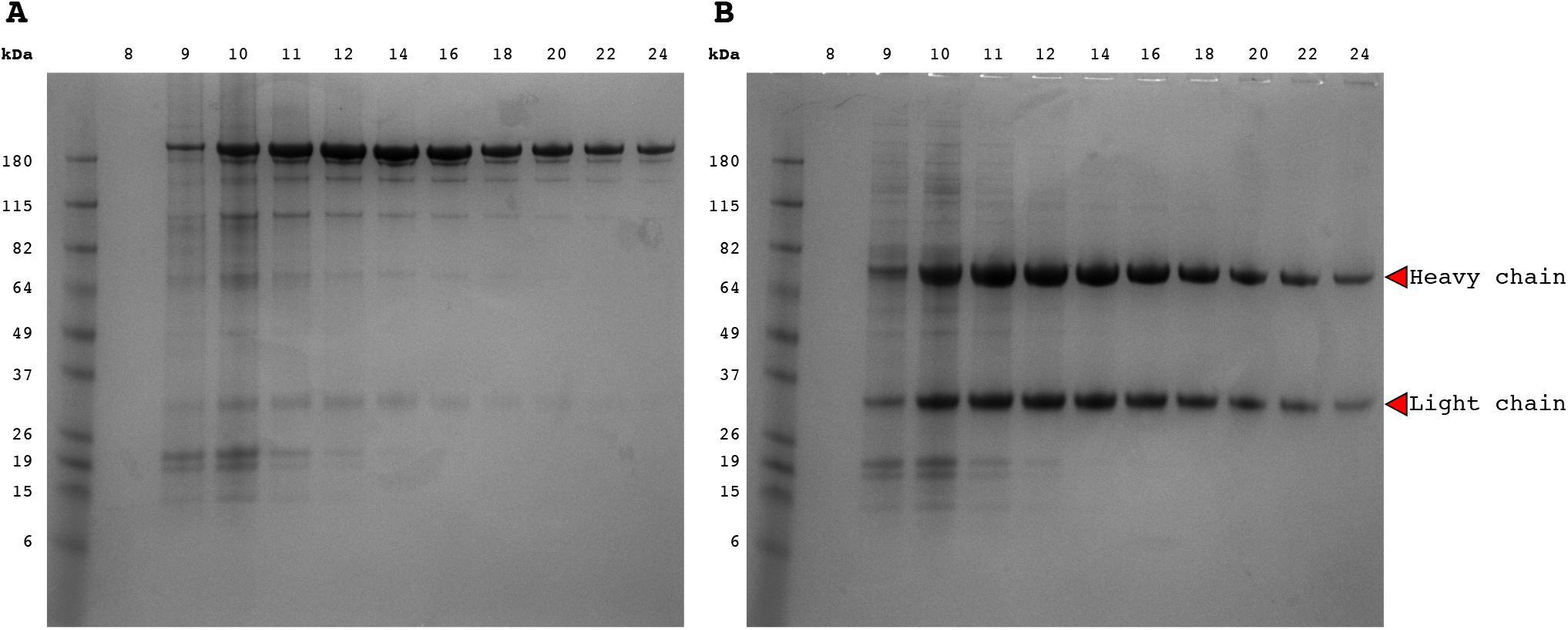
Fractions collected from a step elution (see Fig. 6) in CR3022 mAb purification using an affinity chromatography (5-mL HiTrap™ Protein G HP column). The fractions were analysed on a non-reducing **(A)** and reducing **(B)** SDS-PAGE, confirming the presence of both heavy and light chains.

### HSPG was co-purified with full-length SARS-CoV-2 spike

During our optimization of protein expression in CHO cells, a longer cell cultivation time was found to have a profound and positive effect on the full-length spike yield (data not shown). However, the spike purified from the spent medium of a longer culture did not provide the same purity as seen in Fig. 2A, despite applying the same purification scheme (Fig. 8A). Notably, we observed protein species of much higher Mw (>250 kDa) and of lower Mw (predominantly at ∼70 kDa). Western blot analysis with an anti-His antibody showed that these co-purified proteins were not His-tagged (Fig. 8B), and the possibility of spike degradation due to a longer cell cultivation time was ruled out.

**Figure 8:**
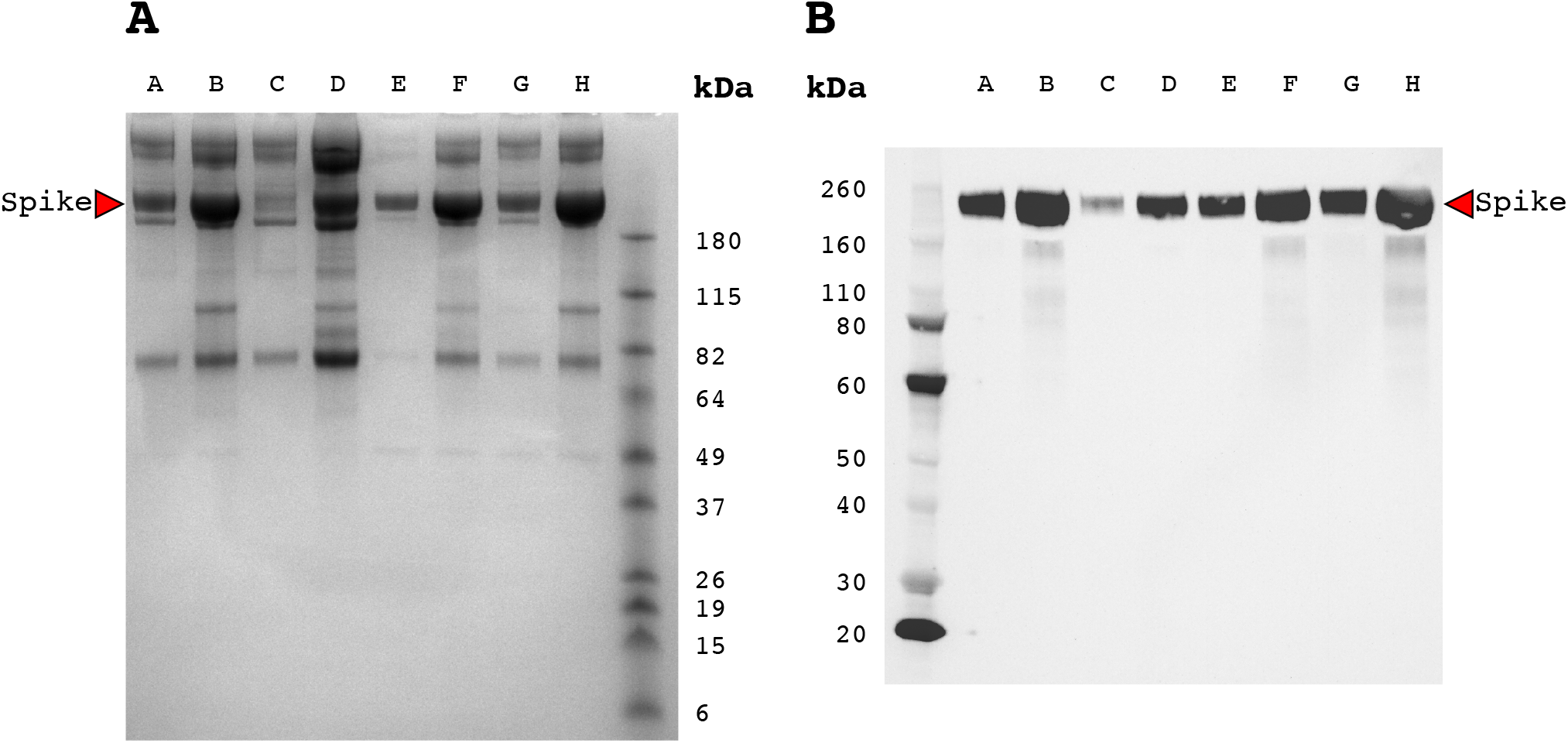
**(A)** Purified spike from a CHO culture with a long cultivation time was less pure compared to purified spike in Fig. 2. Main protein impurities were of higher (>250 kDa) and lower (predominantly at ∼70 kDa) Mw. **(B)** Western blot analysis of the same protein samples in (A) using an anti-His antibody.

To identify these impurities, two impure spike fractions (Fig. 8A, lanes D and F) were subjected to in-solution trypsin digestion and mass spectrometry analysis. The two fractions gave almost an identical protein profile (Table 1). Based on the mass spectrometry data, heparan sulfate proteoglycan (HSPG, Mw 334 kDa), nidogen-1 (79 kDa) and suprabasin (64 kDa) constituted bulk of the co-purified impurities, consistent with the impurity bands (high Mw of >250 kDa and low Mw of ∼70 kDa) observed in the SDS-PAGE analysis (Fig. 8A).

**Table 1:** Protein impurities (Fig. 8, lanes D and F), co-purified with the full-length spike, as determined by mass spectrometric analysis. Proteins are ranked from high to low abundance.

| Impure spike fraction 1 (Fig. 8, lane D) |  |  |  | Impure spike fraction 2 (Fig 8, lane F) |  |  |  |
| --- | --- | --- | --- | --- | --- | --- | --- |
| Protein | UniProt ID | M<sub>w</sub> | %His | Protein | UniProt ID | M<sub>w</sub> | %His |
| Basement membrane-specific heparan sulfate proteoglycan core protein | G3HIM1 | 334 kDa | 4.3% | Basement membrane-specific heparan sulfate proteoglycan core protein | G3HIM1 | 334 kDa | 4.3% |
| Nidogen-1 | G3HWE4 | 79 kDa | 3.0% | Suprabasin | G3HPT8 | 64 kDa | 8.1% |
| Tubulointerstitial nephritis antigen-like | G3H1W4 | 53 kDa | 3.0% | Nidogen-1 | G3HWE4 | 79 kDa | 3.0% |
| Suprabasin | G3HPT8 | 64 kDa | 8.1% | Nidogen-1 | G3I3U5 | 30 kDa | 3.2% |
| Nidogen-1 | G3I3U5 | 30 kDa | 3.2% | Tubulointerstitial nephritis antigen-like | G3H1W4 | 53 kDa | 3.0% |
| Inter-alpha-trypsin inhibitor heavy chain H5 | G3GR64 | 102 kDa | 2.2% | Inter-alpha-trypsin inhibitor heavy chain H5 | G3GR64 | 102 kDa | 2.2% |
| Lysyl oxidase homolog | G3I7U8 | 88 kDa | 4.1% | MAM domain-containing protein 2 | G3IFK8 | 53 kDa | 2.8% |
| Granulins | G3HLK3 | 64 kDa | 5.1% |  |  |  |  |
| Lamin-A/C | G3HG95 | 64 kDa | 1.4% |  |  |  |  |
| Laminin subunit gamma-1 | G3HG25 | 172 kDa | 1.9% |  |  |  |  |

### Peaks corresponding to HSPG and full-length spike were incompletely resolved using a gradient elution in affinity chromatography or a size exclusion chromatography

To further optimise the affinity chromatography step, two strategies were attempted: (a) a gradient elution, and (b) a more stringent binding by adding 20 mM imidazole into the spent medium prior to sample loading.

In Fig. 9, a gradient elution [0 – 100% (v/v) buffer B over 5 CVs] was applied to an impure spike fraction (Fig. 8, lane D). HSPG interacts strongly with Ni sepharose, as the peak corresponding to HSPG was eluted at 47% (v/v) buffer B. Although it was possible to partially resolve the peaks corresponding to HSPG (fractions 13 – 16) and spike (fractions 15 – 24), as shown in Fig. 10A, this elution scheme was not further pursued for two reasons. First, the spike protein was eluted in a broad peak spanning over 10 fractions. This lowers the concentration of the purified spike and would mean a larger volume to process in the subsequent desalting step. The lower spike concentration is not ideal as protein is typically more stable when stored at a higher concentration. Second, some spike protein was lost through co-elution with HSPG (Fig. 10, fractions 15 and 16). Adding 20 mM imidazole into the spent medium, however, provided a clear benefit of higher purity without compromising the protein yield (Fig. 10B).

**Figure 9:**
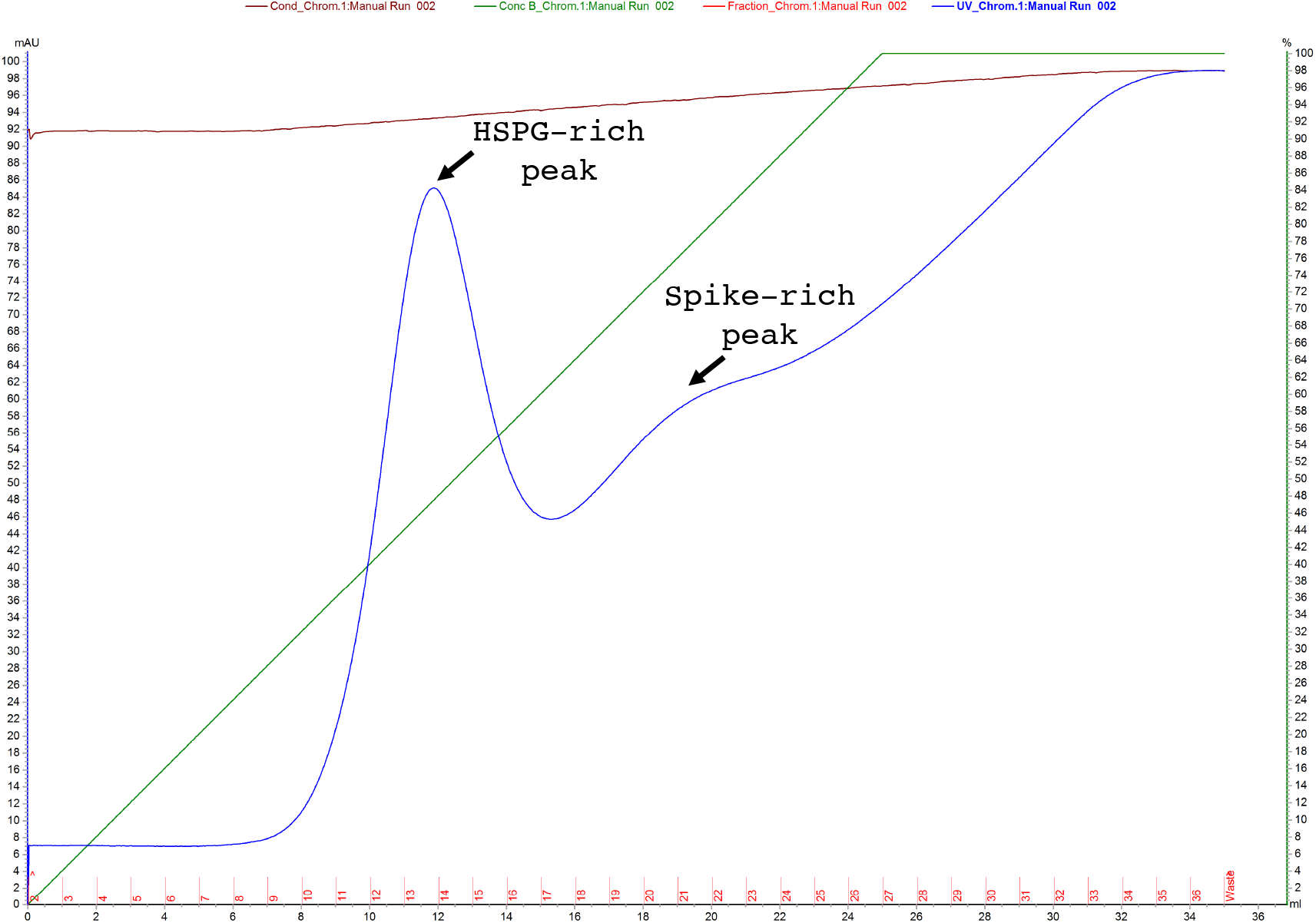
The chromatogram of a gradient elution [0 – 100% (v/v) buffer B over 5 column volumes], when an impure spike fraction (Fig. 8, lane D) was loaded onto a 5-mL HisTrap™ HP column. The x axis, left y axis, and right y axis show the elution volume in mL, UV absorbance at 280 nm, and % (v/v) of buffer B, respectively. The blue, green and brown curves represent UV absorbance at 280 nm, % (v/v) of buffer B and conductivity, respectively. HSPG interacted strongly with Ni sepharose. The peak corresponding to HSPG was eluted at 47% (v/v) buffer B.

**Figure 10:**
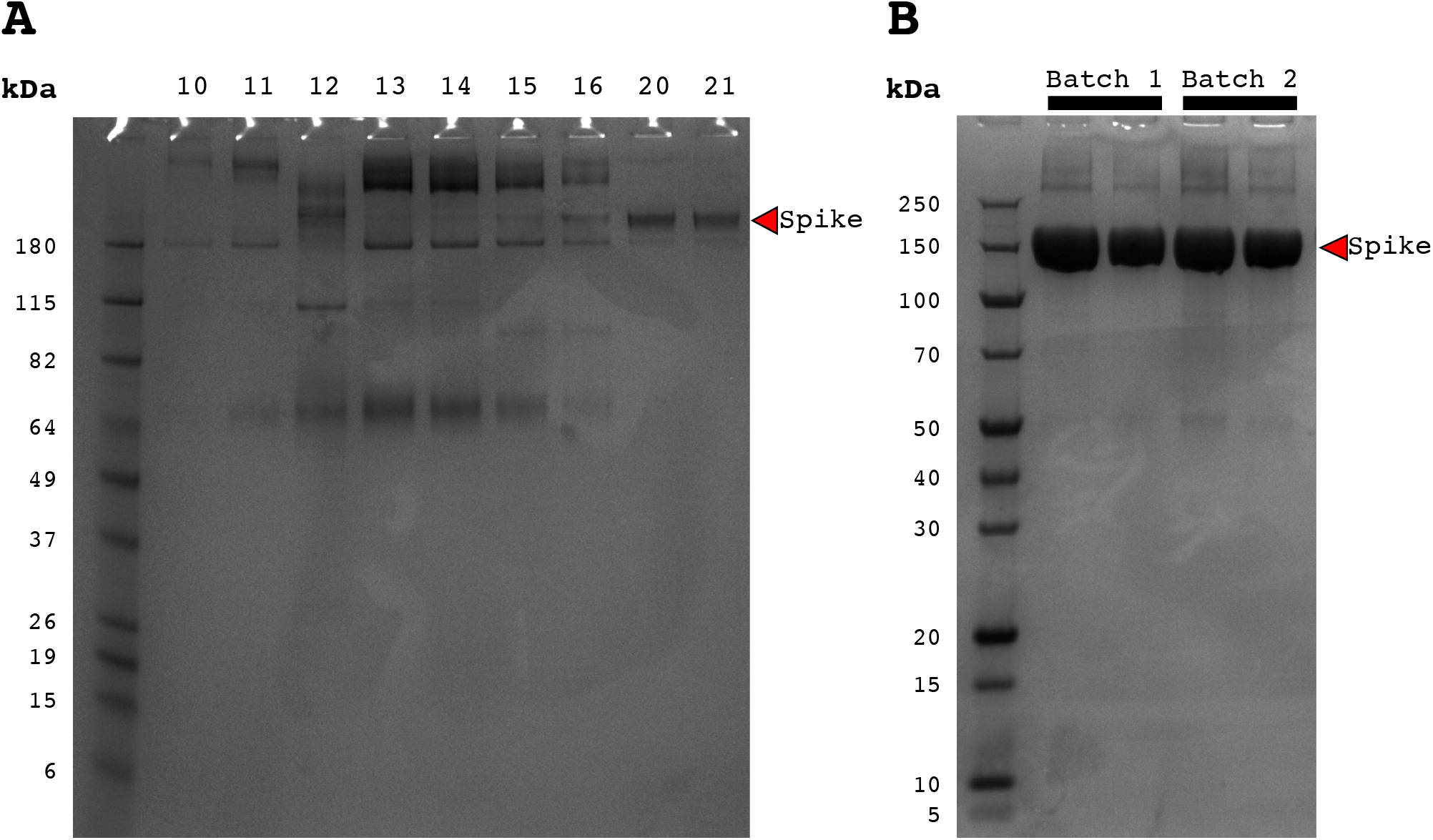
**(A)** Fractions collected from a gradient elution [0 – 100% (v/v) buffer B over 5 column volumes, see Fig. 9], when an impure spike fraction (Fig.8, lane D) was loaded onto a 5-mL HisTrap™ HP column. HSPG (fractions 13 – 16) and spike (fractions 15 – 24) were partially resolved. **(B)** A more stringent binding by adding 20 mM imidazole into the spent medium, prior to sample loading onto a 5-mL HisTrap™ HP column, improved the purity of spike when the protein was purified from CHO culture with a long cultivation time.

We also attempted size exclusion chromatography (SEC) using an impure spike fraction, with the aim of simultaneously achieving (a) separation between HSPG and spike, and (b) buffer exchange. However, we obtained a broad peak (Fig. 11, line 2) with a slight shoulder (line 3). SDS-PAGE analysis confirmed that the peak corresponding to HSPG was eluted at ∼116 mL, while the one corresponding to spike at ∼121 mL (Fig. 12). Although SEC was not the right approach to separate HSPG from spike, we could deduce the Mw of the full-length spike from the calibration curve for a HiLoad™ 26/600 Superdex™ 200 pg column (Fig. 13A). Using an elution volume of ∼121 mL, the recombinant spike has a Mw of 619 kDa (Fig. 13B), confirming the trimeric nature of this glycoprotein. This Mw is in good agreement with the one reported by Watanabe *et al.* using a similar full-length spike construct (670 kDa, deduced from SEC using a Superose 6 10/300 column) [18].

**Figure 11:**
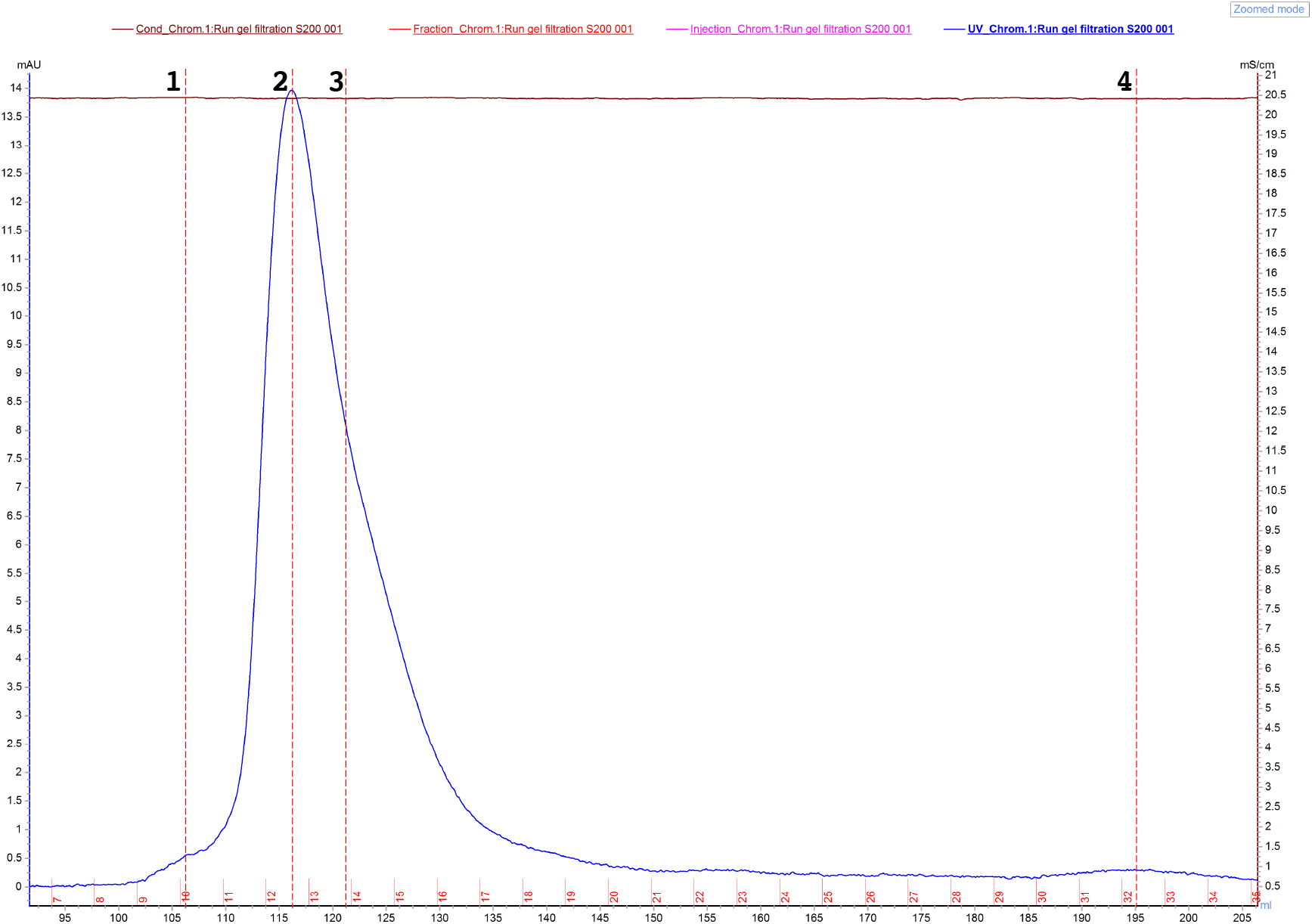
The chromatogram of a size exclusion chromatography, when a 2-mL impure spike fraction (Fig.8, lane D) was loaded onto a HiLoad™ 26/600 Superdex™ 200 pg column. The x axis, left y axis, and right y axis show the elution volume in mL, UV absorbance at 280 nm, and % (v/v) of buffer B, respectively. The blue, green and brown curves represent UV absorbance at 280 nm, % (v/v) of buffer B and conductivity, respectively. Lines 1 – 4 showed the four peaks in the chromatogram.

**Figure 12:**
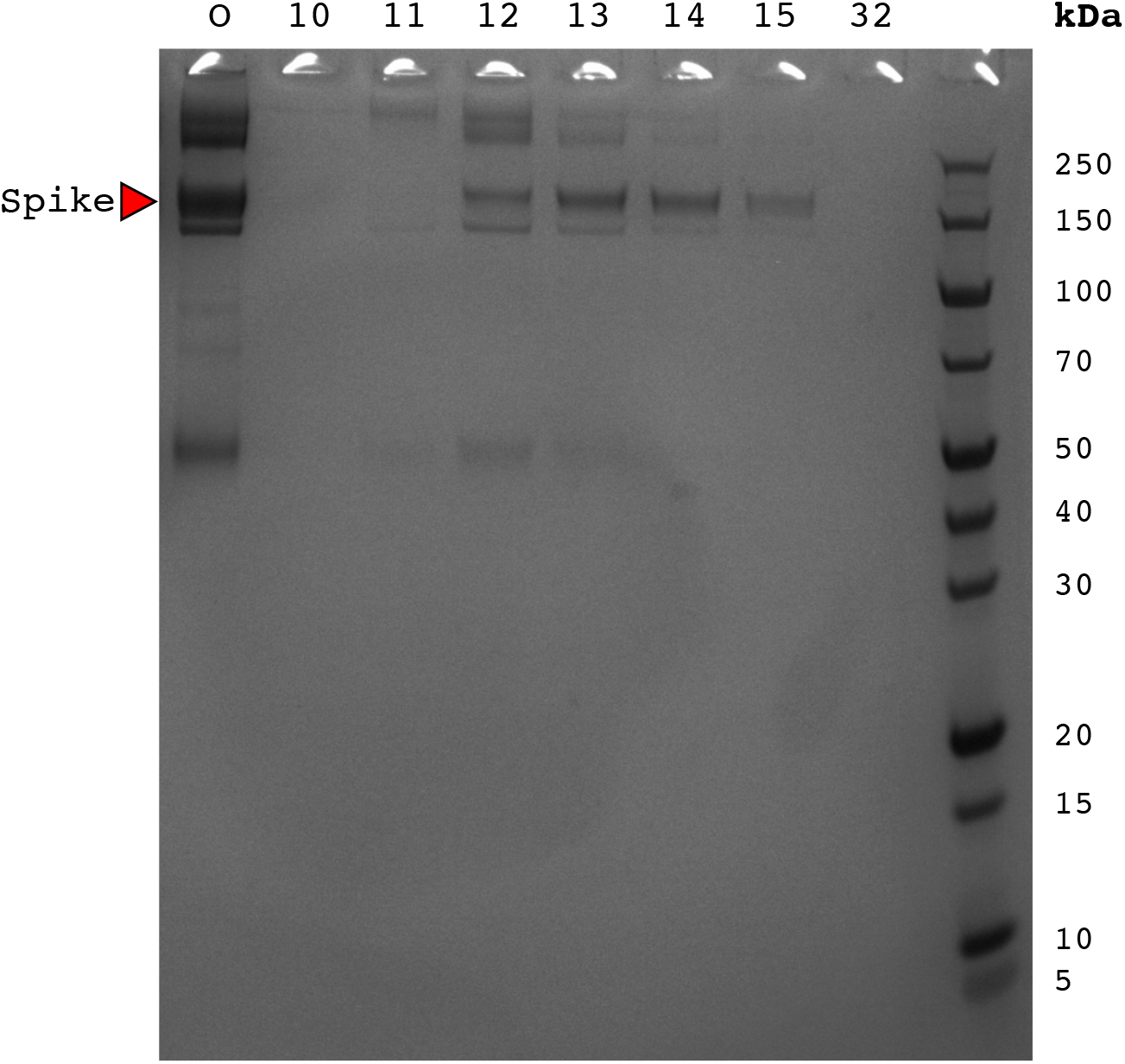
Fractions collected from the size exclusion chromatography (SEC, see Fig. 11), when a 2-mL impure spike fraction (Fig.8, lane D) was loaded onto a HiLoad™ 26/600 Superdex™ 200 pg column. O represents the sample prior to SEC.

**Figure 13:**
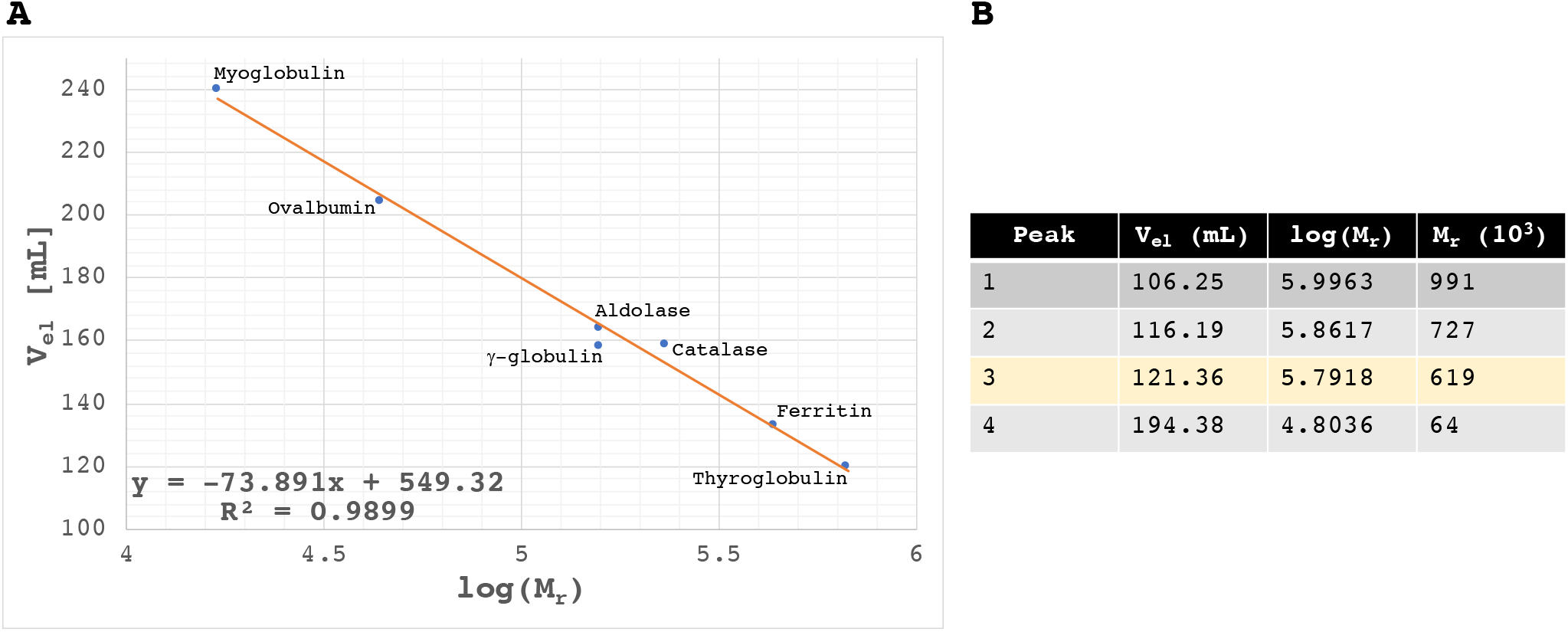
**(A)** The calibration curve for a HiLoad™ 26/600 Superdex™ 200 pg column, created using the data obtained from EMBL’s Protein Expression and Purification Core Facility (https://www.embl.de/pepcore/pepcore_services/protein_purification/chromatography_/hiload26-60_superdex200/). **(B)** Calculated Mw of the four peaks, indicated by lines 1 – 4 in the chromatogram shown in Fig. 11. Spike (line 3 in Fig. 11) has a Mw of 619 kDa, confirming its trimeric nature.

### Co-immunoprecipitation

Co-purification of HSPG raises the possibility that HSPG could potentially be a binding partner of SARS-CoV-2 spike. To further investigate the potential HSPG and spike interaction, four protein samples were prepared and loaded onto NAb™ Protein G spin columns: sample 1 (75 μg CR3022), sample 2 (75 μg CR3022 + 71 μg spike), sample 3 (75 μg CR3022 + 50 μg protein from the HSPG-rich fraction), and sample 4 (75 μg CR3022 + 50 μg protein from the HSPG-rich fraction + 71 μg spike). The HSPG-rich fraction used in these samples referred to the peak corresponding to HSPG from a gradient elution (Fig. 9).

As shown in Fig. 14, all the impurities including HSPG were not captured by the Protein G spin column. They were present in the flowthrough and absent from the eluate. Spike, on the other hand, was captured by the Protein G spin column via CR3022. Although we did not observe a co-complex formed between spike and HSPG, we could not rule out their potential interaction for two reasons. First, co-immunoprecipitation relies on strong affinities between the bait (CR3022) and the primary target (spike) as well as the latter with the secondary target (HSPG). Low affinity or transient interaction between proteins may not be detected. The spike-HSPG interaction could potentially be a weak one. Second, CR3022 could be a competitor of HSPG for spike interaction. Although the co-immunoprecipitation experiment was inconclusive with respect to spike-HSPG interaction, it confirmed nonetheless (a) the functionality of the purified spike and CR3022, and (b) the incomplete resolution of the peaks corresponding to HSPG and spike in a gradient elution. Therefore, if a gradient elution was adopted during purification, the spike protein loss would be higher than the gain in purity.

**Figure 14:**
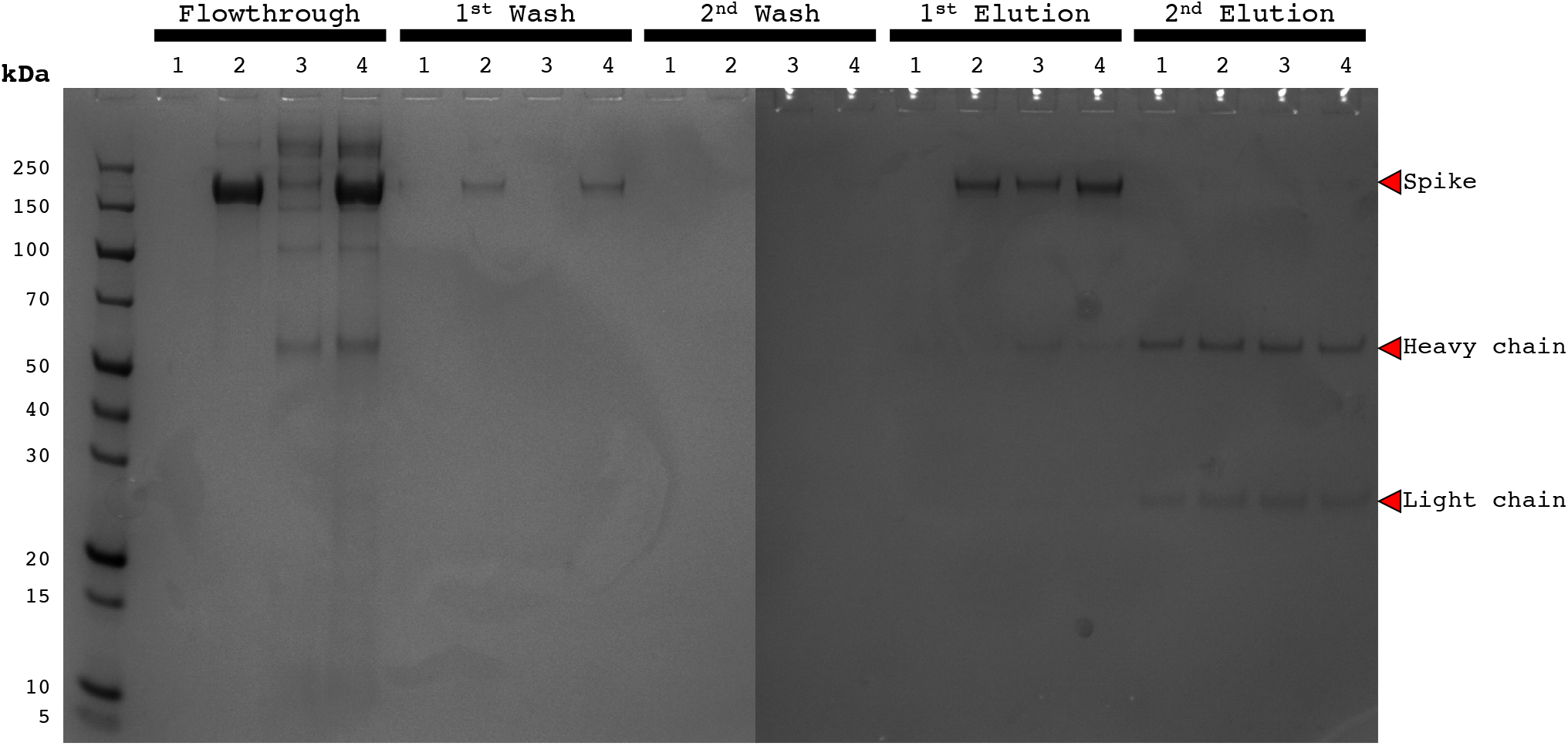
Four protein samples were prepared and loaded onto NAb™ Protein G spin columns: sample 1 (75 μg CR3022), sample 2 (75 μg CR3022 + 71 μg spike), sample 3 (75 μg CR3022 + 50 μg protein from the HSPG-rich fraction), and sample 4 (75 μg CR3022 + 50 μg protein from the HSPG-rich fraction + 71 μg spike). The HSPG-rich fraction used in these samples referred to the peak corresponding to HSPG from a gradient elution (see Fig. 9). The samples from flowthrough, 1^st^ washing, 2^nd^ washing, 1^st^ elution, and 2^nd^ elution were analysed on a reducing SDS-PAGE.

### Co-purification of HSPG

HSPGs are composed of unbranched, negatively charged heparan sulfate polysaccharides attached to a variety of cell surface or extracellular matrix proteins [21]. Owing to the heavily sulfated glycosaminoglycan chains, they present a global negative charge that interacts electrostatically with the basic residues of viral surface glycoproteins or viral capsid proteins. Viruses exploit these weak interactions with HSPG to increase their local concentration at the cell surface, augmenting their chances of binding a more specific receptor [21]. HSPGs are expressed in lung cells [22]. They also play a pivotal role in cellular internalization of viruses, basic peptides and polycation-nucleic-acid complexes [23].

Both heparan sulfate [24] and heparin [25] were recently shown to interact with SARS-CoV-2 spike and its RBD, further strengthening our postulation that CR3022 and HSPG could compete with one another for spike binding. All these evidences in the literature, along with the experimental observation presented in this study, suggest that HSPG co-purification is not a coincidental event. We hypothesize that HSPG copurification is contributed by two synergistic mechanisms, as illustrated in Fig. 15A. First, HSPG interacts with spike via electrostatic interaction. As spike is being captured by the HisTrap™ column, the high local concentration of spike within the column would favour protein-protein interaction. Second, the high histidine content of HSPG (135 residues, 4.3%) further augment its binding with Ni sepharose or spike. The pKa value of the histidine side chain is 6.0. Some of these histidine residues would be deprotonated in our purification buffers of pH 8.0. Important to note, three arginine residues and one lysine residue were removed from the spike variant used in this study (R682, R683, R985 and K986). These modifications would have compromised the affinity of HSPG to spike. Recently, the D614G mutation in spike was shown to increase the infectivity of the virus [26]. Removal of a negatively charged residue from spike could benefit its interaction with HSPG.

**Figure 15:**
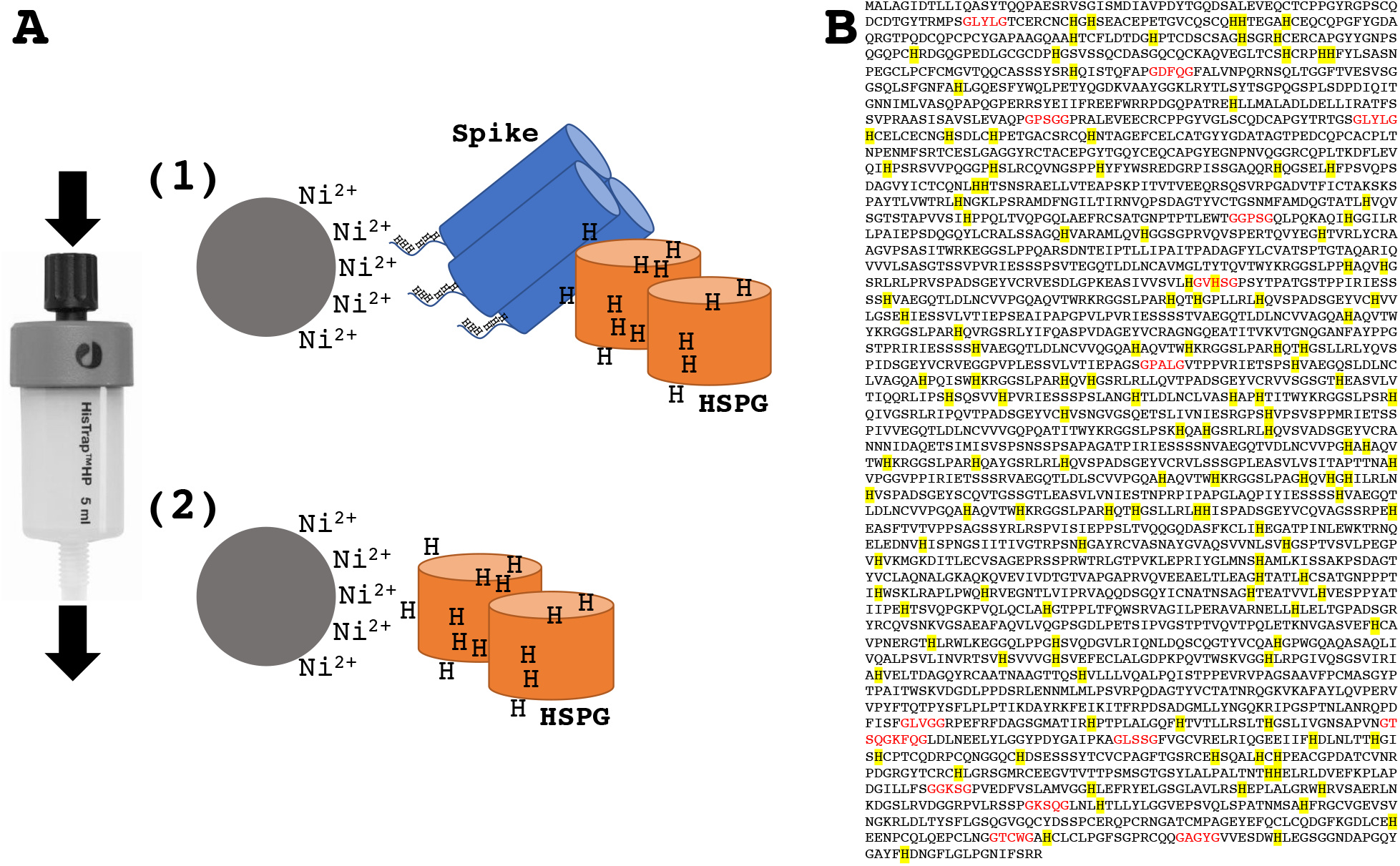
**(A)** HSPG co-purification is hypothetically contributed by two synergistic mechanisms. (1) HSPG interacts with spike via electrostatic interaction. (2) The high histidine content of HSPG (135 residues, 4.3%) further augment its binding with Ni sepharose or spike. **(B)** The protein sequence of HSPG (UnitProt G3HIM1). All histidine residues are highlighted in yellow. GXXXG dimerization motifs are coloured in red.

Based on our SEC data (Fig. 13b), HSPG has a Mw of 727 kDa, in line with a previous study (400 – 600 kDa, purified from mouse cells) [27]. There are a number of GXXXG motifs in the protein sequence of HSPG (Fig. 15B), suggesting that this protein could form a dimer [28].

### Co-purification of proteins other than HSPG

A close examination of other impurities revealed multiple extracellular matrix (ECM) macromolecules, such as laminin, nidogen and tubulointerstitial nephritis antigen-like [29]. They were captured by the Ni sepharose for two reasons: (a) a high histidine content *(e.g.,* 8.1% histidine in suprabasin), and (b) interaction with HSPG. Laminin, an ECM-adhesive protein, is known to interact with HSPG [30]. Nidogen, another impurity we identified, interacts with both laminin [31] and HSPG [32].

Throughout our spike purifications, we have consistently noticed that the spike purified from a HEK culture was purer than that from a CHO culture. This is in agreement with a previous report that CHO cells had a higher amount of cell-surface and intracellular HSPGs compared to HEK293 cells [33].

## CONCLUSION

Recombinant spike, its RBD and CR3022, purified using the purification schemes presented here, are adequate for routine serological assays, both in terms of protein purity and yield. Although CHO is an excellent manufacturing host for spike, impurities such as HSPG are co-purified from CHO culture with a long cultivation time when a HisTrap column is used for spike purification. One potential mitigation strategy is the use of HSPG-deficient CHO cells or those defective in the biosynthesis of glycosaminoglycans [34]. HEK293, with a lower HSPG expression, is a good alternative. The potential HSPG-spike interaction certainly warrants further investigation, as it may contribute to the pathology and the pathogenesis of SARS-CoV-2 and other coronaviruses.

## ACKNOWLEDGEMENT

We thank Prof. Florian Krammer (The Icahn School of Medicine, Mount Sinai) for sharing the SARS-CoV-2 spike and RBD constructs, and acknowledge the support from the Royal Academy of Engineering (the Leverhulme Trust Senior Research Fellowship; to TSW; LTSRF1819\15\21), the University of Sheffield (GCRF Fellowship; to KLT), and the Biotechnology and Biological Sciences Research Council UK (to MJD; BB/M012166/1).

## REFERENCES

1. Zhu, N., et al., A Novel Coronavirus from Patients with Pneumonia in China, 2019. N Engl J Med, 2020. 382(8): p. 727–733.

2. To, K.K., et al., Temporal profiles of viral load in posterior oropharyngeal saliva samples and serum antibody responses during infection by SARS-CoV-2: an observational cohort study. Lancet Infect Dis, 2020. 20(5): p. 565–574.

3. Tortorici, M.A. and D. Veesler, Structural insights into coronavirus entry. Adv Virus Res, 2019. 105: p. 93–116.

4. Shang, J., et al., Structural basis of receptor recognition by SARS-CoV-2. Nature, 2020. 581(7807): p. 221–224.

5. Walls, A.C., et al., Structure, Function, and Antigenicity of the SARS-CoV-2 Spike Glycoprotein. Cell, 2020. 181(2): p. 281–292 e6.

6. Zheng, M. and L. Song, Novel antibody epitopes dominate the antigenicity of spike glycoprotein in SARS-CoV-2 compared to SARS-CoV. Cell Mol Immunol, 2020. 17(5): p. 536–538.

7. ter Meulen, J., et al., Human monoclonal antibody combination against SARS coronavirus: synergy and coverage of escape mutants. PLoS Med, 2006. 3(7): p. e237.

8. Zhou, P., et al., A pneumonia outbreak associated with a new coronavirus of probable bat origin. Nature, 2020. 579(7798): p. 270–273.

9. Tian, X., et al., Potent binding of 2019 novel coronavirus spike protein by a SARS coronavirus-specific human monoclonal antibody. Emerg Microbes Infect, 2020. 9(1): p. 382–385.

10. Yuan, M., et al., A highly conserved cryptic epitope in the receptor binding domains of SARS-CoV-2 and SARS-CoV. Science, 2020. 368(6491): p. 630–633.

11. Amanat, F., et al., A serological assay to detect SARS-CoV-2 seroconversion in humans. Nat Med, 2020.

12. Stadlbauer, D., et al., SARS-CoV-2 Seroconversion in Humans: A Detailed Protocol for a Serological Assay, Antigen Production, and Test Setup. Curr Protoc Microbiol, 2020. 57(1): p. e100.

13. Wu, F., et al., A new coronavirus associated with human respiratory disease in China. Nature, 2020. 579(7798): p. 265–269.

14. Braun, E. and D. Sauter, Furin-mediated protein processing in infectious diseases and cancer. Clin Transl Immunology, 2019. 8(8): p. e1073.

15. Pallesen, J., et al., Immunogenicity and structures of a rationally designed prefusion MERS-CoV spike antigen. Proc Natl Acad Sci U S A, 2017. 114(35): p. E7348–E7357.

16. Guthe, S., et al., Very fast folding and association of a trimerization domain from bacteriophage T4 fibritin. J Mol Biol, 2004. 337(4): p. 905–15.

17. Tao, Y., et al., Structure of bacteriophage T4 fibritin: a segmented coiled coil and the role of the C-terminal domain. Structure, 1997. 5(6): p. 789–98.

18. Watanabe, Y., et al., Site-specific glycan analysis of the SARS-CoV-2 spike. Science, 2020.

19. Lan, J., et al., Structure of the SARS-CoV-2 spike receptor-binding domain bound to the ACE2 receptor. Nature, 2020. 581(7807): p. 215–220.

20. Akerstrom, B. and L. Bjorck, A physicochemical study of protein G, a molecule with unique immunoglobulin G-binding properties. J Biol Chem, 1986. 261(22): p. 10240–7.

21. Cagno, V., et al., Heparan Sulfate Proteoglycans and Viral Attachment: True Receptors or Adaptation Bias? Viruses, 2019. 11(7).

22. Haeger, S.M., Y. Yang, and E.P. Schmidt, Heparan Sulfate in the Developing, Healthy, and Injured Lung. Am J Respir Cell Mol Biol, 2016. 55(1): p. 5–11.

23. Belting, M., Heparan sulfate proteoglycan as a plasma membrane carrier. Trends Biochem Sci, 2003. 28(3): p. 145–51.

24. Liu, L., et al., SARS-CoV-2 spike protein binds heparan sulfate in a length- and sequence-dependent manner. bioRxiv, 2020: p. 2020.05.10.087288.

25. Mycroft-West, C.J., et al., Heparin inhibits cellular invasion by SARS-CoV-2: structural dependence of the interaction of the surface protein (spike) S1 receptor binding domain with heparin. bioRxiv, 2020: p. 2020.04.28.066761.

26. Korber, B., et al., Tracking changes in SARS-CoV-2 Spike: evidence that D614G increases infectivity of the COVID-19 virus. Cell, 2020.

27. Paulsson, M., et al., Structure and function of basement membrane proteoglycans. Ciba Found Symp, 1986. 124: p. 189–203.

28. Yano, Y., et al., GXXXG-Mediated Parallel and Antiparallel Dimerization of Transmembrane Helices and Its Inhibition by Cholesterol: Single-Pair FRET and 2D IR Studies. Angew Chem Int Ed Engl, 2017. 56(7): p. 1756–1759.

29. Theocharis, A.D., D. Manou, and N.K. Karamanos, The extracellular matrix as a multitasking player in disease. FEBS J, 2019. 286(15): p. 2830–2869.

30. Carey, D.J., et al., Association of cell surface heparan sulfate proteoglycans of Schwann cells with extracellular matrix proteins. J Biol Chem, 1990. 265(33): p. 20627–33.

31. Dziadek, M., M. Paulsson, and R. Timpl, Identification and interaction repertoire of large forms of the basement membrane protein nidogen. EMBO J, 1985. 4(10): p. 2513–8.

32. Sage, J., et al., Cleavage of nidogen-1 by cathepsin S impairs its binding to basement membrane partners. PLoS One, 2012. 7(8): p. e43494.

33. Lee, S., et al., Heparan sulfate proteoglycan synthesis in CHO DG44 and HEK293 cells. Biotechnology and Bioprocess Engineering, 2016. 21(3): p. 439–445.

34. Esko, J.D., T.E. Stewart, and W.H. Taylor, Animal cell mutants defective in glycosaminoglycan biosynthesis. Proc Natl Acad Sci U S A, 1985. 82(10): p. 3197–201.

35. Sievers, F., et al., Fast, scalable generation of high-quality protein multiple sequence alignments using Clustal Omega. Mol Syst Biol, 2011. 7: p. 539.

36. Pettersen, E.F., et al., UCSF Chimera--a visualization system for exploratory research and analysis. J Comput Chem, 2004. 25(13): p. 1605–12.

